# The Effects of Global Signal Regression on Estimates of Resting-state BOLD fMRI and EEG Vigilance Correlations

**DOI:** 10.1101/433912

**Authors:** Maryam Falahpour, Alican Nalci, Thomas T. Liu

## Abstract

Global signal regression (GSR) is a commonly used albeit controversial preprocessing approach in the analysis of resting-state BOLD fMRI data. While the effects of GSR on resting-state functional connectivity measures have received much attention, there has been relatively little attention devoted to its effects on studies looking at the relation between resting-state BOLD measures and independent measures of brain activity. In this study we used simultaneously acquired EEG-fMRI data in humans to examine the effects of GSR on the correlation between resting-state BOLD fluctuations and EEG vigilance measures. We show that GSR leads to a positive shift in the correlation between the BOLD and vigilance measures. This shift leads to a reduction in the spatial extent of negative correlations in widespread brain areas, including the visual cortex, but leads to the appearance of positive correlations in other areas, such as the cingulate gyrus. The results obtained using GSR are consistent with those of a temporal censoring process in which the correlation is computed using a temporal subset of the data. Since the data from these retained time points are unaffected by the censoring process, this finding suggests that the positive correlations in cingulate gyrus are not simply an artifact of GSR.

## 1. Introduction

In the past two decades, resting state blood oxygen-level-dependent (BOLD) functional magnetic resonance imaging (fMRI) has become a popular tool for studying the functional organization of the brain. The BOLD signal is a complex mixture of neuronal and non-neuronal components including motion, physiological fluctuations, and thermal noise. Although there have been many advances in methods for removing non-neuronal components (Liu, 2016; Caballero-Gaudes and Reynolds, 2017), there is still no gold standard for the preprocessing of resting-state BOLD fMRI data.

One controversial preprocessing step is the use of the global signal as a nuisance regressor (Liu et al., 2017). The global signal (GS) is defined as the average time series computed over all voxels within the brain and its inclusion as a nuisance regressor is commonly referred to as global signal regression (GSR). Since the GS is by definition a time-varying spatial average, it is useful for representing the influence of spatially coherent noise sources, such as motion and physiological noise (Liu, 2016). Prior studies have shown that GSR can improve the specificity of positive correlations (Fox et al., 2009) in resting-state functional connectivity maps and significantly reduce motion and respiratory signal artifacts (Power et al., 2015). However, Murphy et al. (2009) argued that GSR can introduce artificial negative correlations. Building on these concerns, Saad et al. (2012) further argued that GSR could have a spatially varying bias on connectivity measures, where the bias varies with the true underlying correlations. On the other hand, Nalci et al. (2017) have reported that GSR can be approximated as a temporal downweighting or a censoring function and concluded that anti-correlations observed in the default mode network were not solely an artifact of GSR. More recently, several studies using simultaneous EEG and fMRI measures have demonstrated a relation between the global signal and EEG measures of vigilance (Schölvinck et al., 2010; Wong et al., 2013; Falahpour et al., 2016), indicating the contribution of neural activities to the global signal.

The use of GSR in simultaneous EEG-fMRI studies is less prevalent and it is not always clear if it has been employed. In prior studies where GSR does not appear to have been used, both EEG alpha power and vigilance (defined as the ratio of alpha power to theta and delta power) have been found to be negatively correlated with the BOLD signal in widespread regions of the brain including the lingual gyrus, posterior cingulate, cuneus, and precuneus (Goldman et al., 2002; Laufs et al., 2003; Falahpour et al., 2018). Furthermore, these studies reported that positive EEG-BOLD correlations were primarily observed in the thalamus. In a study which used GSR, Sadaghiani et al. (2010) found positive correlations in additional areas not reported in prior studies, including the dorsal anterior cingulate cortex, the anterior insula, and the anterior prefrontal cortex. We hypothesized that the discrepancy in the findings was partly due to the differences in the preprocessing approaches. To test this hypothesis, we used simultaneously acquired EEG/fMRI data to investigate the correlations between EEG vigilance and BOLD fMRI both prior to and after GSR.

## 2. Methods

### 2.1. Experimental protocol

In this work we used data from the protocol described in our prior work (Wong et al., 2013). Details of the experimental protocol, acquisition and MR/EEG data pre-processing were described in that article and are repeated here for the reader’s convenience. Twelve healthy volunteers were initially enrolled in this study after providing informed consent. Two subjects were not able to complete the entire study, resulting in a final sample size of 10 subjects (4 males and 6 females, Aged 24-33 years, with an average age of 25.6 years). A repeated measures design was used, with each subject participating in two imaging sessions: a caffeine session and a control session. The order of the two sessions was randomized in a double-blinded manner. Each scan section consisted of (1) a high-resolution anatomical scan, (2) arterial spin labeling scans (these results are reported in our prior study (Wong et al., 2012) but not considered here), and (3) two 5 minute resting-state scans with simultaneous EEG recording (one eyes-closed and one eyes-open). Only the eyes-closed scans from non caffeine sessions were used in this work. Subjects were instructed to lie still in the scanner and not fall asleep during resting-state scans. During the eyes-closed (EC) resting-state scans, subjects were asked to imagine a black square. An impedance check was performed prior to each EEG data acquisition while the subject was in the desired state (EC or EO) for at least 1.5 min. EEG data acquisition began 30 s before each fMRI scan and ended 30 s after. Thus, the subject was in the desired state (EC or EO) for at least 2 min prior to the acquisition of the combined EEG and fMRI data. Field maps were acquired to correct for magnetic field inhomogeneities.

### 2.2. MR data acquisition

Imaging data were acquired on a 3 Tesla GE Discovery MR750 whole body system using an eight-channel receiver coil. High resolution anatomical data were collected using a magnetization prepared 3D fast spoiled gradient (FSPGR) sequence (TI = 600 ms, TE = 3.1 ms, flip angle = 8°, slice thickness = 1 mm, FOV = 25.6 cm, matrix size =256 × 256 × 176). Whole brain BOLD resting-state data were acquired over 30 axial slices using an echo planar imaging (EPI) sequence (flip angle = 70°, slice thickness = 4 mm, slice gap = 1 mm, FOV = 24 cm, TE = 30 ms, TR = 1.8 s, matrix size = 64 × 64 × 30). Field maps were acquired using a gradient recalled acquisition in steady state (GRASS) sequence (TE1 = 6.5 ms, TE2 = 8.5 ms), with the same in-plane parameters and slice coverage as the BOLD resting state scans. The phase difference between the two echoes was then used for magnetic field inhomogeneity correction of the BOLD data. Cardiac pulse and respiratory effect data were monitored using a pulse oximeter (InVivo) and a respiratory effort transducer (BIOPAC), respectively. The pulse oximeter was placed on each subject’s right index finger while the respiratory effort belt was placed around each subject’s abdomen. Physiological data were sampled at 40 Hz using a multi-channel data acquisition board (National Instruments).

### 2.3. EEG data acquisition

EEG data were recorded using a 64 channel MR-compatible EEG system (Brain Products, Munich, Germany). The system consisted of two 32 channel Brain Amp MR Plus amplifiers powered by a rechargeable battery unit. The system was placed behind the scanner bore, which was connected using a 125 cm long data cable to a Brain Cap MR with 64 recording electrodes. All electrodes in the cap had sintered Ag/AgCl sensors incorporating 5 *k*Ω safety resistors. The separate ECG electrode had a built-in 15 kΩ resistor. The arrangement of the electrodes in the cap conformed to the international 10/20 standard. FCz and AFz were the reference and ground electrodes, respectively. The EEG data were recorded at a 5*kHz* sampling rate with a passband of 0.1-250 Hz. A phase locking device (Syncbox, Brain Products, Munich, Germany) was used to synchronize the clock of the EEG system with the master clock of the MRI system. Before each scan section, the electrode impedances were set below 20 kΩ, while the impedances of the reference and ground electrodes were set below 10 kΩ. Prior to recording EEG data in each resting-state scan, a snapshot of the electrode impedance values was taken from the computer screen. One EEG dataset was created for each 5-min resting-state scan.

### 2.4. MR data processing

AFNI and FSL were used for MRI data pre-processing (Cox, 1996; Smith et al., 2004; Woolrich et al., 2009). The high resolution anatomical data were skull stripped and segmentation was applied to estimate white matter (WM), gray matter (GM) and cerebral spinal fluid (CSF) partial volume fractions. In each scan section, the anatomical volume was aligned to the middle functional volume of the first resting-state run using AFNI. The anatomical volume in the post-dose scan section was then registered to the pre-dose anatomical volume, and the rotation and shift parameters obtained from this registration were applied to the post-dose functional images. The first 6 TRs (10.8 s) of the BOLD data were discarded to allow magnetization to reach a steady state. A binary brain mask was created using the skull-stripped anatomical data. For each slice, the mask was eroded by two voxels along the border to eliminate voxels at the edge of the brain (Rack-Gomer and Liu, 2012). For each run, nuisance terms were removed from the resting-state BOLD time series through multiple linear regression. These nuisance regressors included: i) linear and quadratic trends, ii) six motion parameters estimated during image co-registration and their first derivatives, iii) RETROICOR (2nd order Fourier series) (Glover et al., 2000) and RVHRCOR (Chang et al., 2009; Chang and Glover, 2009) physiological noise terms calculated from the cardiac and respiratory signals, and iv) the mean BOLD signals calculated from WM and CSF regions and their first respective derivatives, where these regions were defined using partial volume thresholds of 0.99 for each tissue type and morphological erosion of two voxels in each direction to minimize the inclusion of gray matter.

### 2.5. EEG data processing

BrainVision Analyzer 2.0.1 software (Brain Products, Munich, Germany) was used for MR gradient removal using an average pulse artifact template procedure (Allen et al., 2000). An average template was created using the volume-start markers from the MRI system and then subtracted from the individual artifacts. After gradient artifact removal, a low pass filter with a cutoff frequency of 30 Hz was applied to all channels and the processed signals were down-sampled to 250 Hz. Heart beat event markers were created within the Analyzer software. The corrected data and event markers were exported to Matlab 7 (Mathworks, Inc.). EEGLAB (version 9) was used for further pre-processing (Delorme and Makeig, 2004). For each EEG dataset, the ballistocardiogram (BCG) and residual artifacts were removed using a combined optimal basis set and independent component analysis approach (OBS-ICA) (Debener et al., 2007; Niazy et al., 2005). To remove the BCG artifact, the continuous EEG data were divided into epochs based on the heart beat event markers. The epochs were stacked into a matrix configuration and a BCG template was created using the first three principal components calculated from the matrix (Debener et al., 2007). The BCG template was then fitted in a least-squares manner and subtracted from each epoch of the EEG data. After BCG artifact removal, channels that exhibited high impedance values (> 20*k*Ω) or were contaminated with high levels of residual gradient artifact were identified and discarded from further processing. The impedance values were identified using the snapshot of channel impedances acquired before the beginning of each scan. To identify channels contaminated with residual gradient artifact that was not adequately removed using the Analyzer software, the continuous EEG data in each channel were bandpass filtered from 15.5 to 17.5 Hz. This frequency band contained the first harmonic of the gradient artifact centered at 16.7 Hz (slice markers were separated by 60 ms). The root mean square (rms) of the filtered time course was calculated for each channel. A channel was identified as a contaminated channel if the rms value was larger than the median plus 6 times the inter-quartile range (Devore and Farnum, 2005), calculated across all channels except the ECG. On average, each channel was included in 95% of the runs in both the eyes-closed and eyes-open conditions (median = 98%, s.d. = 6%). For further analysis, only the included channels for each run were used. Extended infomax ICA was then performed. Independent components (ICs) corresponding to residual BCG or eye blinking artifacts were identified by correlating all IC topographies with artifact template topographies and extracting the ICs with spatial correlation values of 0.8 or more (Debener et al., 2007; Viola et al., 2009). The corrected data were then created by projecting out the artifactual ICs (Delorme and Makeig, 2004). As bulk head motion creates high amplitude distortion in the EEG data acquired in the MRI environment, it is desirable to discard the distorted time segments (Jansen et al., 2012; Laufs et al., 2008). To identify the contaminated time segments, the EEG data were bandpass filtered from 1 to 15 Hz (to avoid the first harmonic of the residual gradient artifact centered at 16.7 Hz). A mean amplitude time series was calculated by taking the rms across channels. Outlier detection was performed on the mean time series and the outlier threshold was calculated as the sum of the median value and 6 times the inter-quartile range across time (Devore and Farnum, 2005). Data points with values larger than this threshold were regarded as segments contaminated by motion. Contaminated segments less than 5 s apart were merged to form larger segments that were then excluded. A binary time series was created by assigning a 1 to the time points within the bad time segments and a 0 to the remaining good time points. In a recent study, (Jansen et al., 2012) found that predictors derived from an EEG motion artifact were strongly correlated with the BOLD signal when the EEG predictors were convolved with a hemodynamic response function. To indicate the time points reflecting the BOLD response to the motion artifact, we convolved the binary time series with a hemodynamic response function (Buxton et al., 2004) and binarized the resulting output with a threshold of zero to form a second binary time series. Since both EEG and fMRI measures were considered in our study, a final binary time series was created by combining the two binary time series using an OR operation. The binary time series were down-sampled to match the sampling frequency of the BOLD time courses. In the eyes-closed condition, an average of 81% of the time points of the binary time series were indicated as good (i.e. with a value of zero; median = 85%, s.d. = 13%). In the eyes-open condition, the corresponding average was 86% (median = 88%, s.d. = 9%). For each EEG channel, a spectrogram was created using a short-time Fourier transform with a 4-term BlackmanHarris window (1311 points, 5.24 s temporal width, and 65.7% overlap). The output temporal resolution of the spectrogram was 1.8 s (i.e. equivalent to the TR of the BOLD resting-state scans). The time points in the EEG spectrogram and BOLD time series that were indicated as potentially contaminated by motion (i.e. values of 1 in the binary time series) were discarded from further analysis.

## 3. FMRI and EEG metrics

### 3.1. EEG vigilance time series

As described above, a spectrogram was calculated for each EEG channel with the same temporal resolution as the BOLD time series. A global spectrogram was created by taking the rms of all spectra across all channels. Then at each time point, relative amplitude spectrum was computed by normalizing the spectrum by its overall rms amplitude (square root of sum of squares across all frequency bins). Relative EEG amplitudes time series were then computed as the rms amplitude in the following frequency bands (delta and theta: 1-7 Hz, alpha: 7-13 Hz). Vigilance defined as the ratio of power in alpha band over the power in delta and theta band is used in the literature (Horovitz et al., 2008; Olbrich et al., 2009; Wong et al., 2013), which is equivalent to the alpha slow-wave index (ASI) (Jobert et al., 1994; Larson-Prior et al., 2009; Müller et al., 2006). In order to form the vigilance time series, rms amplitude in alpha band was divided to the rms amplitude in delta and theta band at each time point. Vigilance time series was then convolved with the hemodynamic response function to account for the hemodynamic delay.

### 3.2. FMRI global signal (GS)

For each voxel, a percent change BOLD time series was obtained by subtracting the mean value from the original voxel time series and then dividing the resulting difference by the mean value. The global signal was formed by averaging the percent change time series across all brain voxels.

### 3.3. Correlation between Vigilance and BOLD

For each run, temporal correlations between the vigilance and fMRI BOLD time courses were first computed before performing GSR. Correlation maps where then normalized using the Fisher-Z transformation. A one sample t-test (3dttest++ from AFNI with t-test randomization to minimize the FPR) was performed on the Fisher-z transformed correlation maps to assess the relation at the group level. The same procedure was repeated after regressing out the GS from the BOLD and vigilance time series.

We further considered a modified implementation of the censoring approach proposed by (Nalci et al., 2017). For each scan, we computed the interquartile range (IQR: middle 50%) of the GS time course and identified the time points for which the GS was outside this range. We then censored the fMRI data from these time points, such that the correlation between vigilance and BOLD was computed using only those time points for which the GS was within the interquartile range. By definition, this approach censors 50 percent of the time points i.e. points below the first quartile (*Q*_1_) or above the third quartile (*Q*_3_).

## 4. Results

### 4.1. BOLD-vigilance correlation before and after GSR

Group result z-score maps showing areas with significant correlation between EEG vigilance and BOLD fMRI are shown in the first and second rows of Figure 1 for data before and after GSR, respectively. Prior to GSR, we found significant positive BOLD-vigilance correlations in the thalamus and significant negative correlations in widespread regions of the brain, including the posterior cingulate, cuneus, precuneus, lingual gyrus, and insula. These findings are consistent with prior studies (Goldman et al., 2002; Laufs et al., 2006) that do not appear to have used GSR. After GSR, there was a significant reduction in the magnitude and the extent of the negative correlations, whereas the extent of the positive correlations in sub-regions of the cingulate gyrus increased. The positive values in the thalamus were found to be largely unaffected. The bottom row in Figure 1 shows z-score maps obtained after censoring based on GS magnitude, which are similar in appearance to the maps obtained after GSR. Supplementary Figure S1 displays the GS time series from a representative subject. The points in the interquartile range (*IQR*), which are used to compute the correlation after censoring, are shown with black circles.

**Figure 1:**
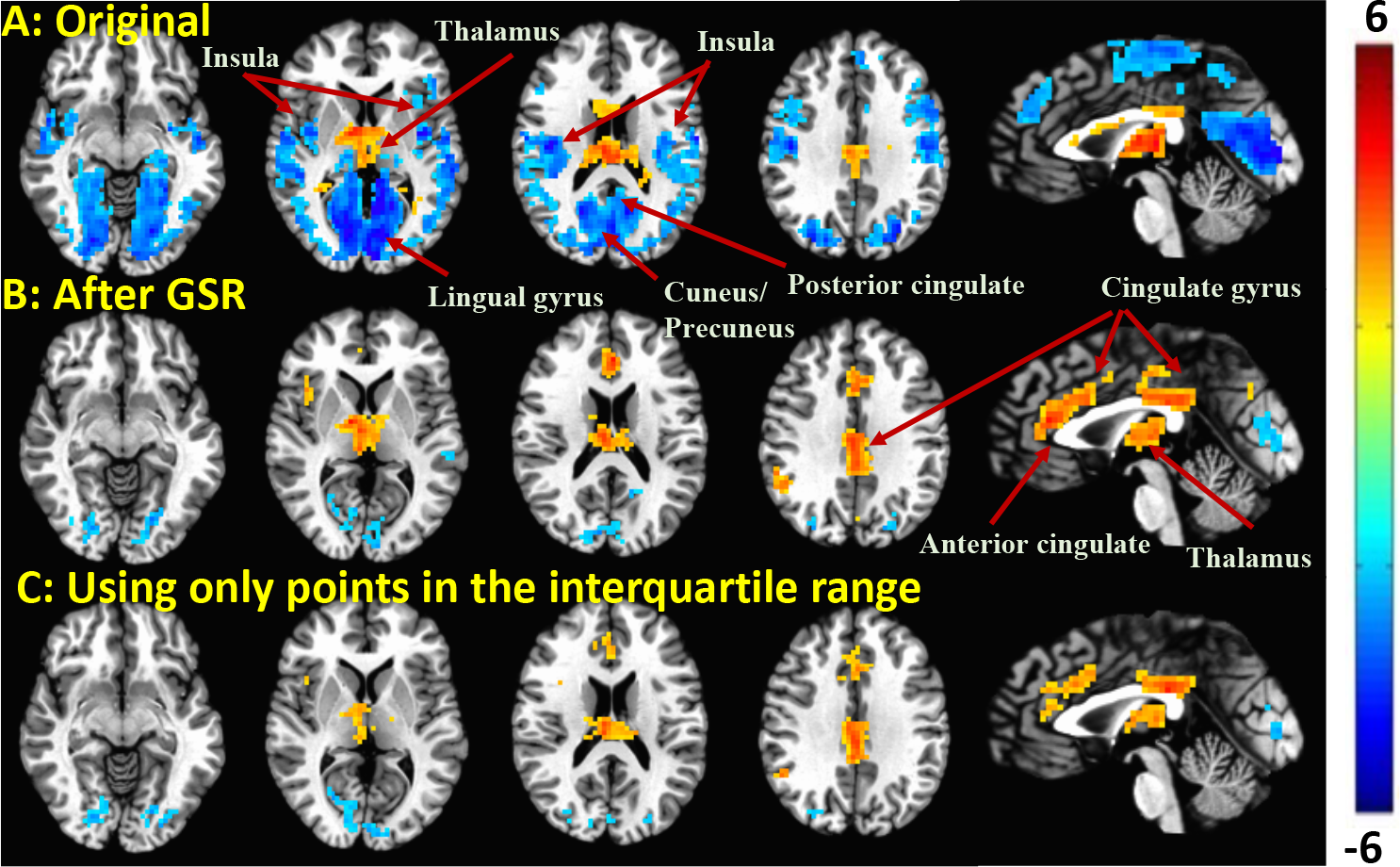
Group result z-scores (from 3ttest++) showing areas of significant correlation between the BOLD fMRI and EEG vigilance when using: A: Original data (before GSR); B: fMRI data after GSR; C: Using only data points in the interquartile range (e.g. marked with black *o* in Figure S1) of the GS. Maps are thresholded at *p* < 0.05 (corrected)

### 4.2. Understanding the changes in correlation values

To gain further insight into the results, we use the Yule expression for partial correlation (Yule, 1897) to write the BOLD-vigilance correlation after GSR as

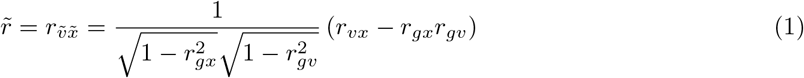

where *x* and *v* denote the BOLD and vigilance time courses prior to GSR, 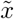 and 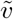 denote the time courses after GSR, *r_vx_* and 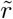 denote BOLD-vigilance correlations before and after GSR, respectively, *r*_*gx*_ and *r*_*gv*_ denote the correlations of the GS with the BOLD and vigilance time courses, respectively. For each voxel, GSR first subtracts an offset term *r*_*gx*_*r*_*gv*_ from the original correlation value. It then multiplies the resulting difference by a scaling term 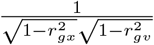, which is by definition greater than or equal to 1.

The offset term *r*_*gx*_*r*_*gv*_ is the product of the (1) correlation *r*_*gv*_ between the GS and vigilance signals and (2) the correlation *r*_*gx*_ between the GS and voxel time series. As noted in our prior work (Falahpour et al., 2018), the GS and vigilance signals exhibit a negative correlation for the vast majority of scans examined (see also Figure 4). On the other hand, the GS is positively correlated with the majority of the voxel time courses in the brain (Fox et al., 2009), reflecting its construction as the average of the voxel time series. In our data, 75% of the voxel time courses were found to be positively correlated with the GS. Taking into account the average behavior of the individual component terms, the offset term *r*_*gx*_*r*_*gv*_ will tend to be negative over the voxels and scans in the study sample, corresponding to a positive shift (due to subtraction of this term) in the correlation values after GSR.

Figure 2 shows an example of the relationship presented in Equation 1 using data from a representative scan. In this example, the scaling term 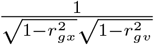 ranges from 1.07 to 2.1. The offset term *r*_*gx*_*r*_*gv*_ consists of a component *r*_*gx*_ that varies across voxels and a component *r*_*gv*_ that is constant across voxels. For this example, *r*_*gv*_ = −0.37 and *r*_*gx*_ is largely positive across voxels, such that the product *r*_*gx*_*r*_*gv*_ is largely negative across voxels. In addition, due to the significant correlation between the GS and vigilance, the spatial structure of the *r*_*gx*_ component is similar to that of the original correlation map *r*_*vx*_. As a result, subtraction of the *r*_*gx*_*r*_*gv*_ term removes much of the spatial structure in the correlation map.

**Figure 2:**
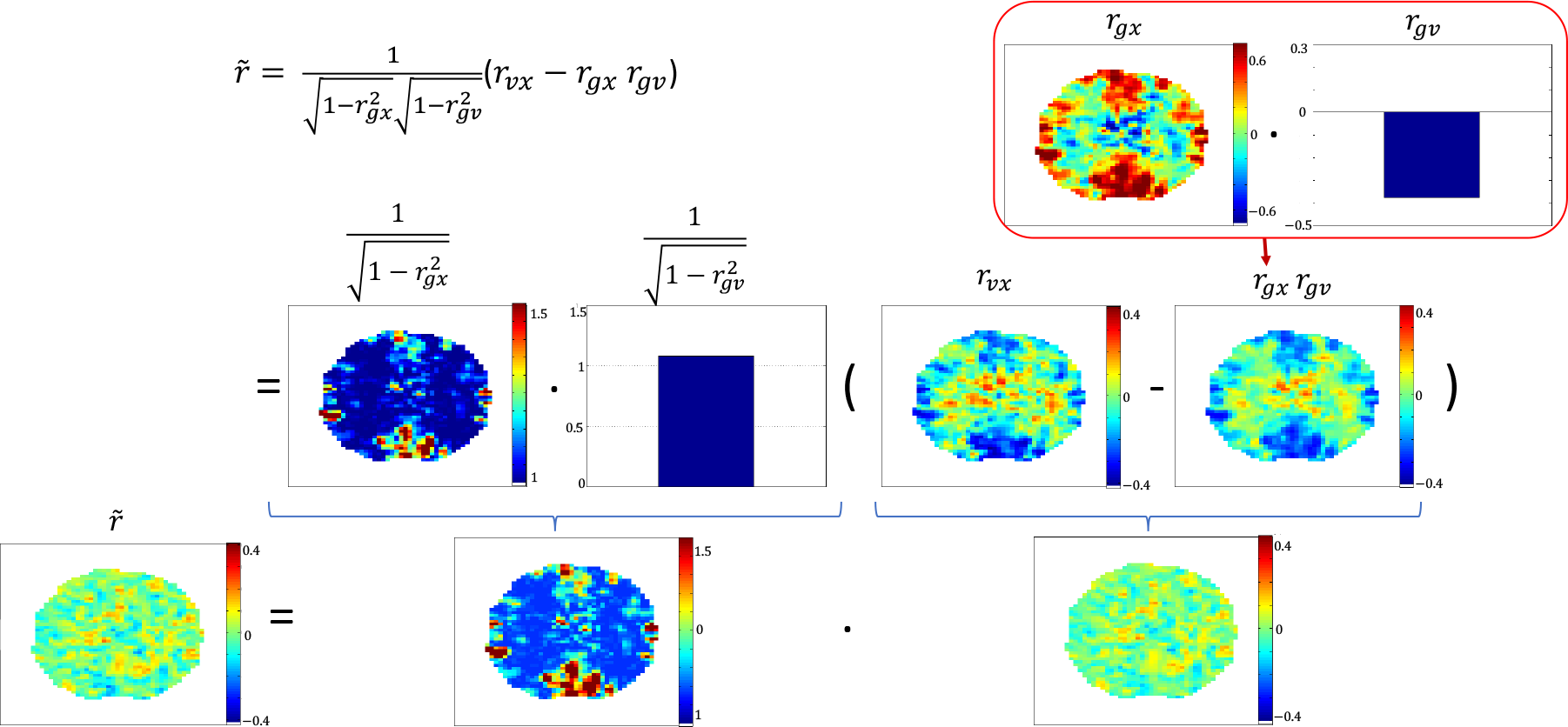
The analytical relation between the BOLD-vigilance correlations before and after GSR is shown graphically for a representative scan. Equation 1 is displayed on the top left side where *v* is the vigilance time series, *x* is the BOLD fMRI time series, *r*_*vx*_ and 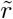 are the correlation between vigilance time series and BOLD fMRI time series before and after GSR, respectively, *r*_*gx*_ is the correlation between GS and BOLD, and *r*_*gv*_ is the correlation between GS and vigilance.

### 4.3. Effect on correlation is primarily explained by subtraction of offset term

In the representative scan, the subtraction of the offset term appears to account for the primary effect of GSR on the BOLD-vigilance correlation map. To assess the relative contributions of the offset and scaling terms across the sample, we computed on a per-scan basis the spatial correlation between the following pairs of maps: (1) BOLD-vigilance correlation map before and after multiplication by the scaling term (first row in Figure 3); (2) BOLD-vigilance correlation map before and after subtraction of the offset term (second row); and (3) BOLD-vigilance correlation map before and after GSR (third row). The spatial correlations shown in the first row are all very close to unity, indicating that multiplication by the scaling term does not significantly change the spatial structure of the correlation maps. In contrast, the similarity of the spatial correlations in the second and third rows indicates that the subtraction of the offset term largely accounts for the change in the spatial structure of the correlation maps after GSR. As further evidence, we show in the fourth row of Figure 3 that there is a high spatial correlation between the BOLD-vigilance maps obtained after GSR and the maps after subtraction of the offset term.

**Figure 3:**
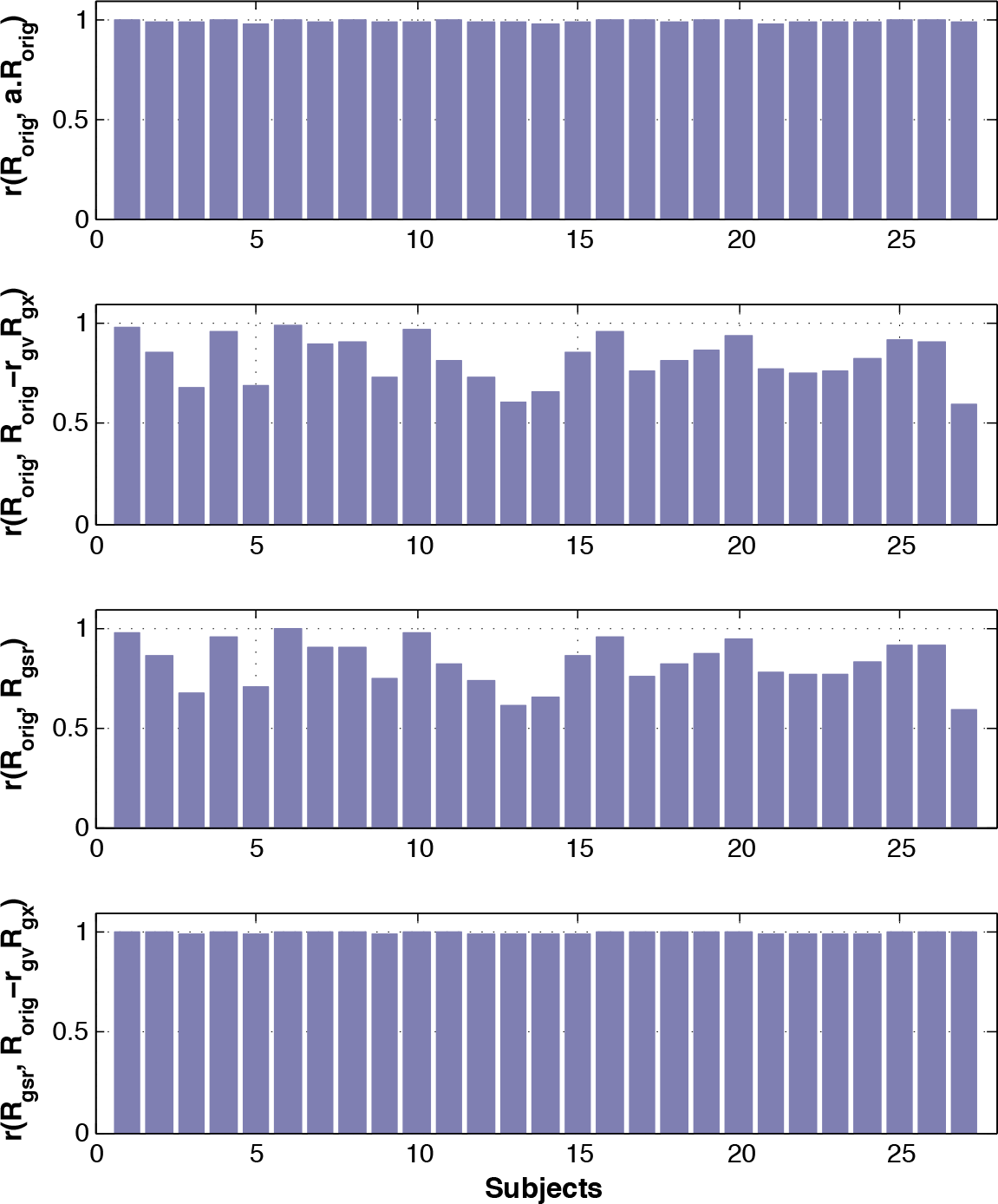
First row: Spatial correlation between BOLD-vigilance correlation maps before and after scaling. Second row: Spatial correlation between BOLD-vigilance correlation maps before and after subtraction of offset term. Third row: Spatial correlation between BOLD-vigilance correlation maps before and after GSR. Fourth row: Spatial correlation between BOLD-vigilance correlation maps obtained after GSR and maps after subtraction of offset term.

### ROI-based analysis

We next examined in detail the changes in correlation values in brain regions where GSR resulted in (1) a decrease in the extent of negative correlations (occipital cortex); (2) an increase in the extent of positive correlations (cingulate gyrus), and (3) no significant changes (thalamus). For analysis, the ROIs were defined using anatomical parcellations from the `TT_caez_ml_18+tlrc` atlas in AFNI as follows: the occipital cortex ROI included Talairach regions 43-48, the cingulate gyrus ROI included Talairach regions 31-32, and the thalamus ROI included Talairach regions 77-78.

Figure 4 displays the terms from Equation 1 for each ROI and scan. In the first row, the original correlation values *r*_*vx*_ are shown, with predominantly negative values for the occipital ROI, largely positive values for the thalamus ROI, and a mix of negative and positive values for the cingulate gyrus ROI. In the second row, the correlation *r*_*gv*_ between the GS and vigilance is shown to be mostly negative across scans (note that these values are independent of the ROI). The correlations *r*_*gx*_ between the GS and ROI time series are shown in the third row. These are uniformly positive for the occipital and cingulate gyrus ROIs, but exhibit a mix of negative and positive values for the thalamus ROI, reflecting a weaker contribution of the thalamic signals to the GS. In the fourth row, we display the product terms *r*_*gx*_*r*_*gv*_ along with the original correlation values *r*_*vx*_ to facilitate comparisons. We find that the product term is largely negative for the occipital and cingulate gyrus ROIs, consistent with the predominantly positive values for *r*_*gx*_ and largely negative values for *r*_*gv*_. In contrast, the product term is roughly centered about zero for the thalamic ROI. The difference *r*_*vx*_ − *r*_*gx*_*r*_*gv*_ between the original correlation and the product term is shown in the bottom row, with scans exhibiting a change in the sign with respect to the original correlation indicated with black asterisks. The cingulate gyrus ROI exhibited the greatest number of sign changes, reflecting the fact that the original values *r*_*vx*_ are roughly centered around zero whereas the subtraction of the offset term causes a largely positive shift. In contrast, the occipital ROI has only a few sign changes because the original values *r*_*vx*_ are predominantly negative and the positive shift due to subtraction of *r*_*gx*_*r*_*gv*_ is not typically sufficient to change the sign. Thus, the correlation values remain negative, but are closer to zero due to the positive shift. Finally, the thalamic ROI displays only 2 sign changes, reflecting the the fact that the original values *r*_*vx*_ are largely positive while the product terms *r*_*gx*_*r*_*gv*_ are rather small in amplitude and centered about zero, such that the resultant changes are relatively small.

**Figure 4:**
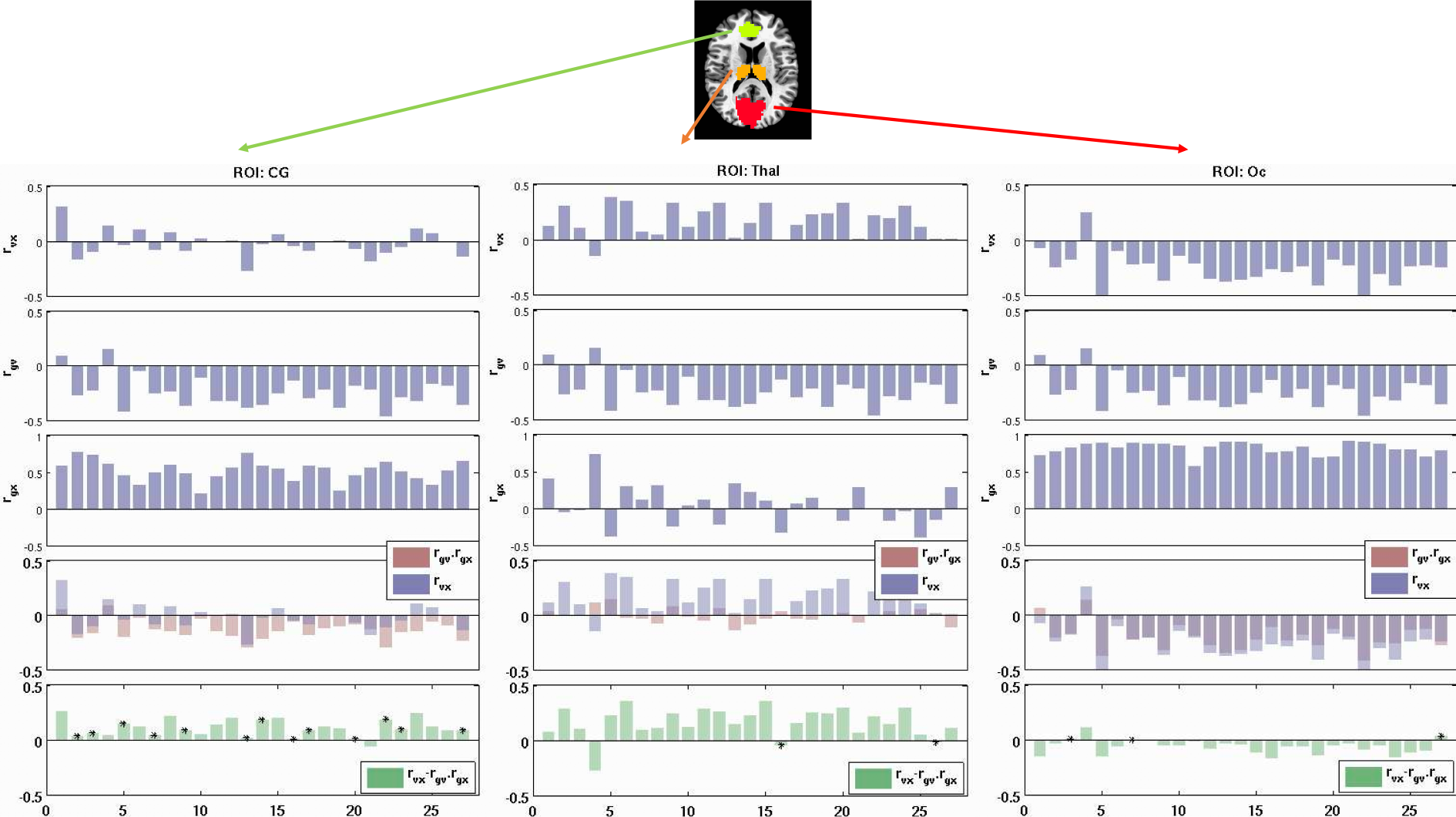
Terms in Equation 1 shown for 3 ROIs and each scan. Columns 1-3 show the results for Cingulate Gyrus (CG), Thalamus (Thal) and the Occipital (Oc) ROIs, respectively. Row1: correlation *r*_*vx*_ between vigilance and the ROI average BOLD signals; Row2: correlation *r*_*gv*_ between GS and vigilance; Row3: correlation *r*_*gx*_ between GS and the ROI average BOLD signals; Row4: multiplication of *r*_*gx*_ and *r*_*gv*_ in pink along with the *r*_*vx*_ in blue; Row5: *r_gx_.r_gv_* subtracted from *r*_*vx*_. Black asterisks show the scans where the polarity of the subtraction is different from the original BOLD-vigilance correlation *r*_*vx*_.

## 5. Discussion

We have shown that GSR alters the correlation between BOLD fluctuations and EEG vigilance measures. After GSR, the correlation maps changed extensively, with a reduction in the extent of a negative correlations in areas such as the visual cortex and an increase in the extent of positive correlations in the cingulate gyrus. These findings are consistent with those of prior studies in which widespread negative correlations between BOLD and alpha power are observed when GSR is not used (Goldman et al., 2002; Laufs et al., 2003; Falahpour et al., 2018) and positive correlations in areas covering the cingulate gyrus are observed when GSR is used (Sadaghiani et al., 2010).

We used the Yule expression for partial correlation (Yule, 1897) to better understand the effects of GSR and examined the relative importance of the scaling and offset subtraction operations. We found that the change in the spatial structure of the correlation maps was primarily due to the subtraction of the spatially varying offset term *r*_*gx*_*r*_*gv*_. Due to the negative correlation *r*_*gv*_ between the GS and vigilance and the largely positive correlation *r*_*gx*_ between the GS and voxel-time series, subtraction of the negative offset term leads to a largely positive shift in the correlation values. In contrast, prior studies have noted that GSR leads to a largely negative shift in correlation values in seed-based functional connectivity maps. For these studies, the term *r*_*gv*_ would be replaced by the correlation between the GS and the seed signal of interest (e.g. *r_gS_* where *S* denotes the seed signal). Since the seed signals used for most functional connectivity maps are positively correlated with the GS, the term *r_gS_* is positive and the subtraction of the corresponding positive offset term *r_gx_r_gS_* leads to a largely negative shift in correlation values. Thus, the difference in the sign of the shifts arises from the differences in the signs of *r*_*gv*_ and *r_gS_*.

We found that the effect of GSR on BOLD-vigilance correlations was spatially varying with pronounced effects in the occipital cortex and cingulate gyrus and relatively little effect in the thalamus. The weak effect in the thalamus reflected the relatively low correlation between the GS and the BOLD signals in that region. In contrast, the higher correlations between the GS and BOLD signals in the both the occipital cortex and cingulate gyrus resulted in a positive shift in the correlation values in these regions. The largely negative correlation values observed before GSR in the occipital cortex were shifted closer to zero by GSR. In contrast, the small positive and negative correlation values before GSR in the cingulate gyrus became largely positive after GSR.

We also found that a censoring approach (in which time points with large GS magnitude are not included in the computation of the correlation maps) provided results comparable to GSR, in line with the findings of (Nalci et al., 2017). This finding suggests that the positive correlations between BOLD and vigilance observed in the cingulate gyrus may not simply be a mathematical artifact introduced by GSR. Note that with the censoring approach there is no mathematical constraint that forces the existence of positive correlations as the images retained for the computation of the correlation coefficient are not modified by censoring. Instead the positive correlations observed in the cingulate gyrus are inherent in the data when considering those time points for which the GS magnitude is relatively low. In addition, the widespread negative BOLD-vigilance correlations are not observed in this temporal subset of the data.

Overall, our observations suggest that negative BOLD-vigilance correlations may largely reflect global fluctuations in BOLD activity and their association with vigilance fluctuations. The reduction of these global fluctuations with GSR leads to an attenuation of the negative BOLD-vigilance correlations in the occipital cortex and reveals positive BOLD-vigilance correlations in the cingulate gyrus. As these positive correlations exist within temporal subsets of the original data, they are unlikely to be caused solely by GSR. Instead, it is possible that these positive correlations are always present but are simply obscured during temporal periods where the GS is high. Future studies are needed to elucidate the mechanisms that give rise to BOLD-vigilance correlations and to better understand why both positive and negative correlations are observed.

---

**Figure S1:**
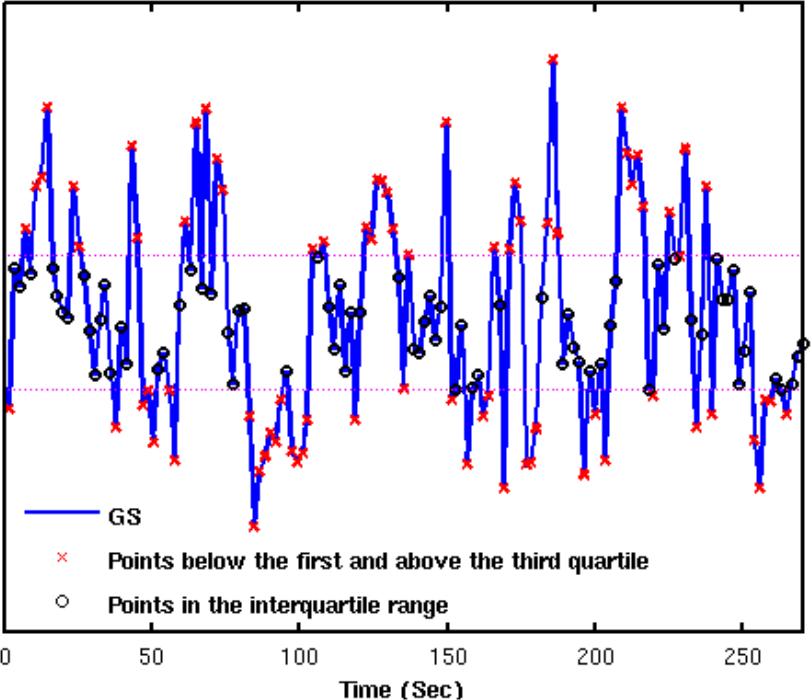
Global signal time series from a representative subject. The points in the interquartile range are marked with with black “o”. Dashed purple lines are Q1 and Q3.

